# Genetic determinants of gut microbiota composition and bile acid profiles in mice

**DOI:** 10.1101/571075

**Authors:** Julia H. Kemis, Vanessa Linke, Kelsey L. Barrett, Frederick J. Boehm, Lindsay L. Traeger, Mark P. Keller, Mary E. Rabaglia, Kathryn L. Schueler, Donald S. Stapleton, Daniel M. Gatti, Gary A. Churchill, Daniel Amador-Noguez, Jason D. Russell, Brian S. Yandell, Karl W. Broman, Joshua J. Coon, Alan D. Attie, Federico E. Rey

## Abstract

The microbial communities that inhabit the distal gut of humans and other mammals exhibit large inter-individual variation. While host genetics is a known factor that influences gut microbiota composition, the mechanisms underlying this variation remain largely unknown. Bile acids (BAs) are hormones that are produced by the host and chemically modified by gut bacteria. BAs serve as environmental cues and nutrients to microbes, but they can also have antibacterial effects. We hypothesized that host genetic variation in BA metabolism and homeostasis influence gut microbiota composition. To address this, we used the Diversity Outbred (DO) stock, a population of genetically distinct mice derived from eight founder strains. We characterized the fecal microbiota composition and plasma and cecal BA profiles from 400 DO mice maintained on a high-fat high-sucrose diet for ∼22 weeks. Using quantitative trait locus (QTL) analysis, we identified several genomic regions associated with variations in both bacterial and BA profiles. Notably, we found overlapping QTL for *Turicibacter sp.* and plasma cholic acid, which mapped to a locus containing the gene for the ileal bile acid transporter, *Slc10a2*. Mediation analysis and subsequent follow-up validation experiments suggest that differences in *Slc10a2* gene expression associated with the different strains influences levels of both traits and revealed novel interactions between *Turicibacter* and BAs. This work illustrates how systems genetics can be utilized to generate testable hypotheses and provide insight into host-microbe interactions.

**Author summary:** Inter-individual variation in the composition of the intestinal microbiota can in part be attributed to host genetics. However, the specific genes and genetic variants underlying differences in the microbiota remain largely unknown. To address this, we profiled the fecal microbiota composition of 400 genetically distinct mice, for which genotypic data is available. We identified many loci of the mouse genome associated with changes in abundance of bacterial taxa. One of these loci is also associated with changes in the abundance of plasma bile acids—metabolites generated by the host that influence both microbiota composition and host physiology. Follow up validation experiments provide mechanistic insights linking host genetic differences, with changes in ileum gene expression, bile acid-bacteria interactions and bile acid homeostasis. Together, this work demonstrates how genetic approaches can be used to generate testable hypothesis to yield novel insight into how host genetics shape gut microbiota composition.

## Introduction

The intestinal microbiota has profound effects on host physiology and health (1–3). The composition of the gut microbiota is governed by a combination of environmental factors, including diet, drugs, maternal seeding, cohabitation, and host genetics (4–7). Together, these factors cause substantial inter-individual variation in microbiota composition and modulate disease risk (8,9). Alterations in the composition of the microbiota are associated with a spectrum of cognitive, inflammatory and metabolic disorders (10–12) and a number of bacterial taxa have been causally linked with modulation of disease (13–15). A major challenge in the field is deciphering how host genetics and environmental factors interact to shape the composition of the gut microbiota. This knowledge is key for designing strategies aimed at modifying gut microbiota composition to improve health outcomes.

Several mouse and human studies have examined the role of host genetics in shaping the composition of the gut microbiota (16). Mouse studies comparing gut bacterial communities from inbred mouse strains (17,18) and strains harboring mutations in immune-related genes (19–22) support this notion. Additionally, quantitative trait locus (QTL) analyses in mice have identified genetic regions associated with the abundance of several bacterial taxa and community structure (23–26). Twin studies and genome-wide association studies (GWAS) in humans have identified heritable bacterial taxa and SNPs associated with specific gut microbes. While comparing these studies is often difficult due to differences in environmental variables among populations, some associations are consistently detected among geographically discrete populations, such as the association between *Bifidobacterium* abundance and the lactase (*LCT*) gene locus (27–29), indicating the abundance of specific taxa is influenced by host genetic variation.

Gut microbes and the host communicate through the production and modification of metabolites, many of which impact host physiology (30–34). Bile Acids (BAs) are host-derived and microbial-modified metabolites that regulate both the gut microbiome and host metabolism (35–37). BAs are synthesized in the liver from cholesterol, stored in the gallbladder and are secreted in the proximal small intestine where they facilitate absorption of fat-soluble vitamins and lipids. Once in the intestine, BAs can be metabolized by gut bacteria through different reactions, including deconjugation, dehydroxylation, epimerization, and dehydrogenation, to produce secondary BAs with differential effects on the host (33,35). In addition to their direct effects on the host, BAs shape the gut microbiota composition through antimicrobial activities (38,39). The detergent properties of BAs cause plasma membrane damage. The bactericidal activity of a BA molecule corresponds to its hydrophobicity (40). Additionally, the microbiota modulates primary BA synthesis through regulation of the nuclear factor FXR (41). Thus, we hypothesized that host genetic variation associated with changes in BA homeostasis mediates alterations in gut microbiota composition.

To investigate how genetic variation affects gut microbiota and BA profiles, we used the Diversity Outbred (DO) mouse population, which is a heterogenous population derived from eight founder strains: C57BL6/J (B6), A/J (A/J), 1291/SvImJ (129), NOD/ShiLtJ (NOD), NZO/HiLtJ (NZO), CAST/EiJ (CAST), PWK/PhJ (PWK), and WSB/EiJ (WSB) (42,43). These eight strains capture a large breadth of the genetic diversity found in inbred mouse strains. Additionally, the founder strains harbor distinct gut microbial communities and exhibit disparate metabolic responses to diet-induced metabolic disease (18,44,45). The DO population is maintained by an outbreeding strategy aimed at maximizing the heterozygosity of the outbred stock. The genetic diversity and large number of generations of outbreeding make it an ideal resource for high-resolution genetic mapping of microbial and metabolic traits (43).

We characterized the intestinal microbiota composition and plasma and cecal BA profiles in ∼400 genetically distinct DO mice fed a high-fat/high-sucrose diet for ∼22 weeks and performed quantitative trait loci (QTL) analysis to identify host genetic loci associated with these traits. Specifically, we focused our analysis on potentially pleiotropic loci, which we defined as a single genetic locus that associates with both bacterial and BA traits. Our analysis revealed several instances of bacterial and metabolite traits attributed to the same DO founder haplotypes mapping to the same position of the mouse genome, including a locus associated with plasma BA levels and the disease-modulating organism *Akkermansia muciniphila.* Additionally, we identified the ileal BA transporter *Slc10a2* as a candidate gene that regulates both the abundance of *Turicibacter sp.* and plasma levels of cholic acid.

## Results and discussion

### Phenotypic variation among Diversity Outbred (DO) mice fed high-fat and high-sucrose diet

We investigated the impact of genetic variation on gut microbiota composition and bile acid (BA) profiles using a cohort of ∼400 DO mice maintained on a high-fat high-sucrose diet (45% kcal from fat and 34% from sucrose) for ∼22 weeks (range 21-25 weeks), starting at weaning. Additionally, we incorporated in our analyses previously published clinical weight traits collected from the same mice (46) (Fig 1A). All animals were individually housed throughout the duration of the study to minimize microbial exchange.

**Figure 1.**
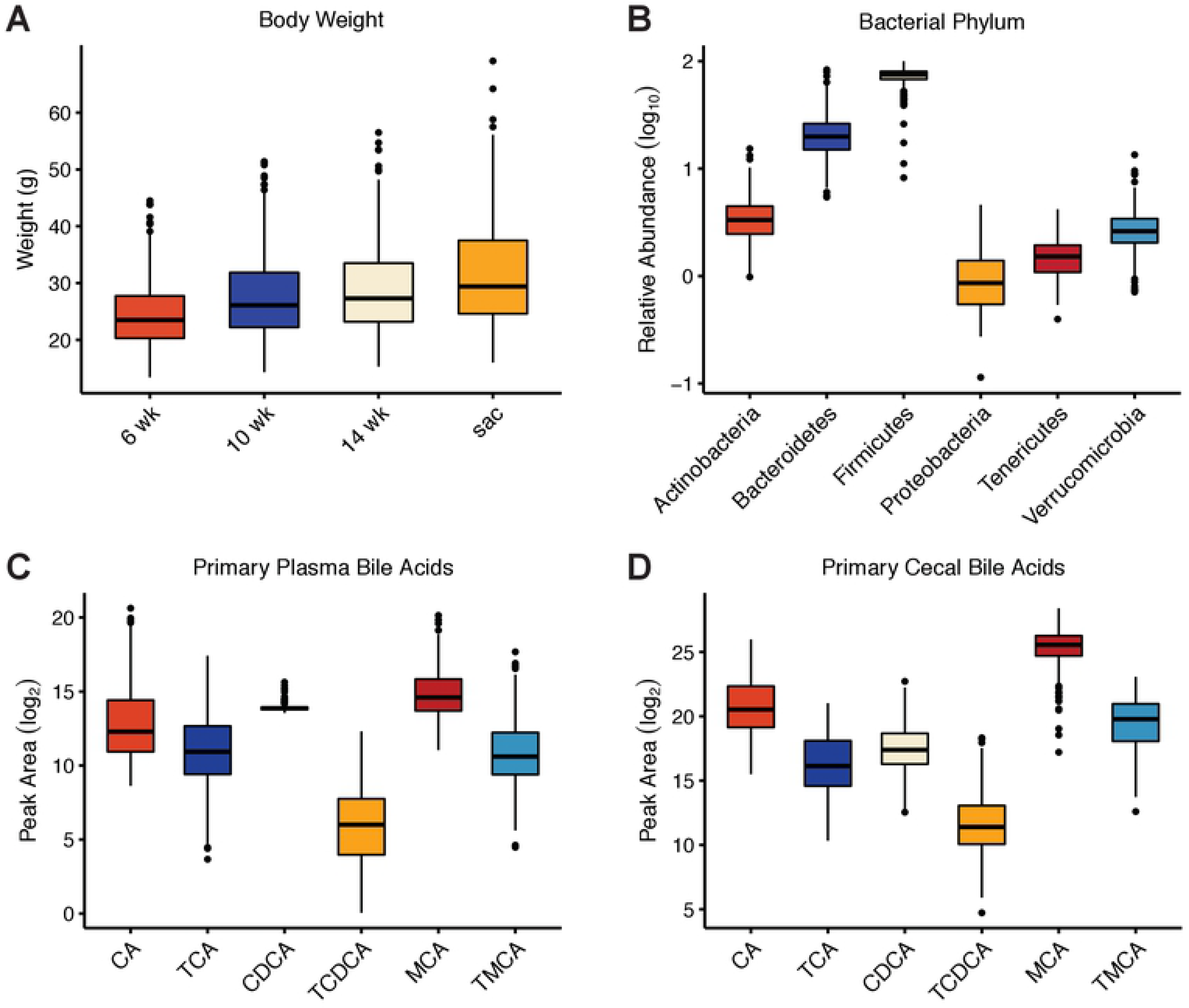
Phenotypic variation among Diversity Outbred (DO) mice fed high-fat and high-sucrose diet. (A) Body weight at 6, 10, 14, and 21-25 (sacrifice) weeks in DO mice fed high-fat and high-sucrose diet (n = 500) (Adapted from Keller et al. (46)) (B) Distributions of the normalized relative abundance of bacterial phyla identified in DO fecal microbiota (n = 399). (C) Abundance (peak area) of primary bile acids detected in plasma and (D) cecal contents (n = 384).

We performed LC-MS analyses of plasma and cecal contents to assess abundance of 27 BAs. There was substantial variation in the plasma and cecal BA profiles across the 400 mice (Fig 1C and 1D; S1 Table). Additionally, we examined gut microbiota composition using 16S rRNA gene amplicon sequencing of DNA extracted from fecal samples collected at the end of the experiment. Within the cohort, there were 907 unique Exact Sequence Variants (ESVs), (100% operational taxonomic units defined with dada2 (47)), which were agglomerated into 151 lower taxonomic rankings (genus, family, order, class, phyla) (S1 Table). The microbial traits represented each of the major phyla found in the intestine and the relative abundance of these phyla was highly variable among the DO mice (Fig 1B). For instance, the abundance of taxa classified to the Bacteroidetes phylum ranged from 1.17 – 89.28%.

For subsequent analysis, we identified a core measurable microbiota (CMM), which we defined as taxon found in at least 20% of the mice (24). This was done to remove the effects of excessive variation in the data due to bacterial taxa that were low abundance and/or sparsely distributed. In total, the CMM was comprised of 86 ESVs and 42 agglomerated taxa (S2 Table). The CMM traits represent a small fraction of the total microbes detected, but account for 94.5% of the rarefied sequence reads, and therefore constitute a significant portion of the identifiable microbiota.

Since mice were received in waves of 100, we examined whether animals in each wave were more similar to each other than mice in other waves. The fecal microbiota composition significantly clustered by wave (p < 0.001, PERMANOVA) and sex (p < 0.001, PERMANOVA) (S1 Fig). PCA analysis of plasma and cecal bile acids showed a significant effect of sex, but not wave, on both plasma (p < 0.0001, Kruskal Wallis) and cecal BA profiles (p < 0.05, Kruskal Wallis) (S2 Fig).

There is substantial evidence implicating gut microbiota and BAs in metabolic disease development (36,37). To identify potential relationships among these traits, we performed correlation analysis which yielded many significant associations after FDR correction (FDR < 0.05) (S3 Table, discussed in S1 Data).

### Abundance of gut bacterial taxa and bile acids are associated with host genetics

To identify associations between regions of the mouse genome and the clinical and molecular traits discussed above, we performed QTL analysis using the R/qtl2 package (48). We used sex, days on the diet, and experimental cohort (wave) as covariates. We identified 459 QTL for bacterial (306), bile acid (131), and body weight (22) traits (Fig 2, S4 Table) with a LOD score > 5.5.

**Figure 2.**
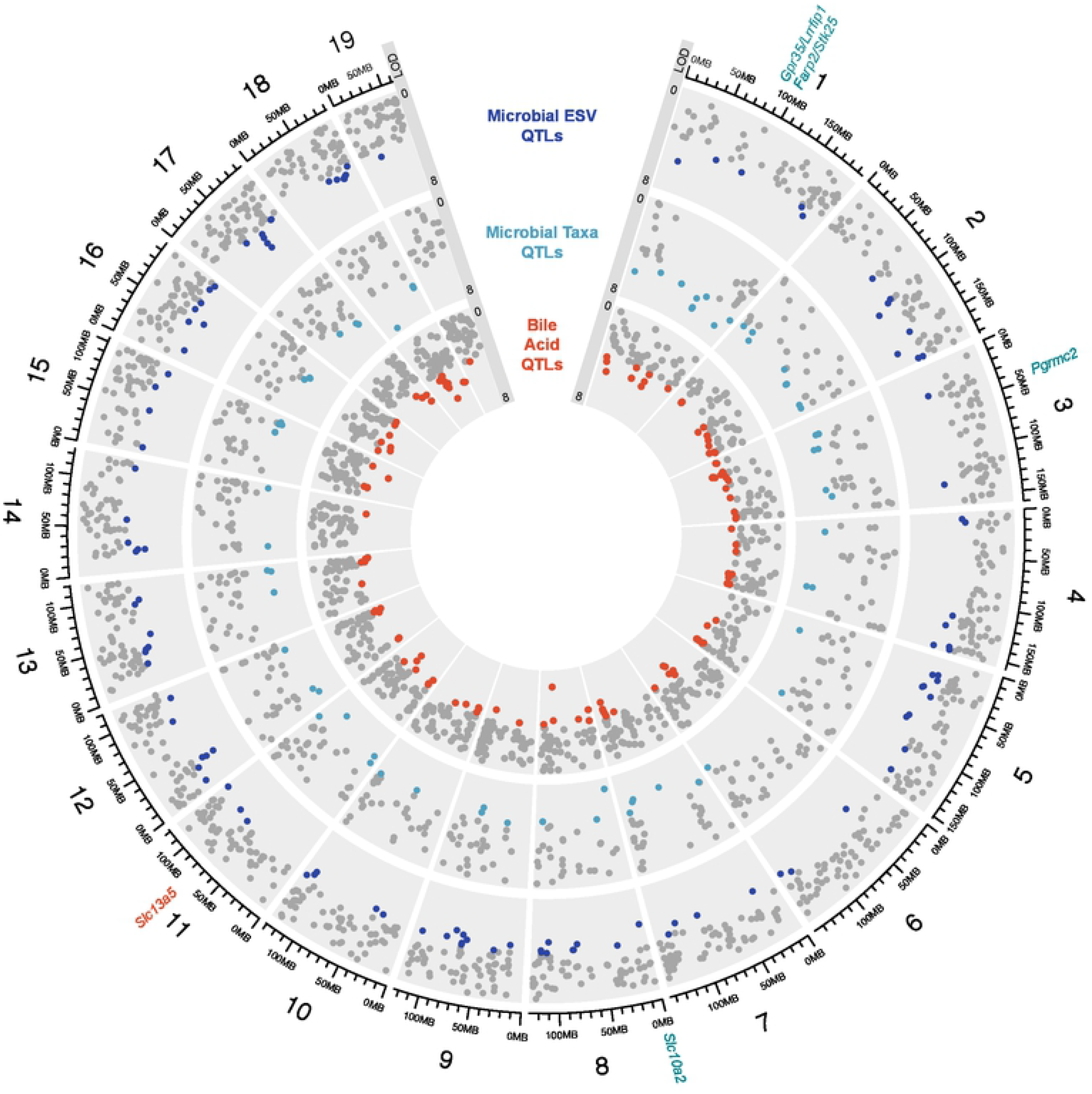
Genetic architecture of quantitative trait loci (QTL) for microbial exact sequence variants (ESVs) and taxa abundance, and plasma and cecal bile acids in 400 Diversity Outbred (DO) mice. The outer layer shows the chromosome location where major tick marks correspond to 25 Mbp. Logarithm of the odds (LOD) range is shown for each track. Each dot represents a QTL on each chromosome of the mouse genome for a given trait. Grey dots denote QTLs with LOD < 5.5. Candidate genes discussed in text are denoted.

Of the microbial QTL, we found 190 QTL for 76 distinct bacterial ESVs from four phyla that met a cut-off LOD > 5.5. ESVs with the strongest QTL (LOD > 8) are classified to the Clostridiales order and map on chr 12 at ∼33 Mbp, the Lachnospiraceae family on chr 2 at 164 Mbp, and the S24-7 family on chr 2 at ∼115 Mbp. We also identified 116 QTL for microbial taxa collapsed by taxonomic assignment (i.e., genus to phylum). The genera *Lactococcus* and *Akkermansia* were also associated with host genetic variation, which is consistent with previous studies (23,24,49,50).

Similarly, BA QTL mapped to multiple loci spanning the mouse genome and most BA traits mapped to multiple positions. BA synthesis and metabolism are regulated by multiple host signaling pathways: there are >17 known host enzymes involved in the production of BAs (36), transporters, which play a critical role in maintaining the enterohepatic circulation and BA homeostasis, and receptors that respond to BA in a variety of host tissues (51–53). Therefore, it is not surprising that our results indicate that BA levels are polygenic and shaped by multiple host factors.

We observed multiple instances of related BA species associating to the same genetic locus. These overlapping QTL may indicate the presence of a pleiotropic locus. Interestingly, several of these loci associate with levels of related BA species in different stages of microbial modification. For example, cecal taurocholic acid (TCA) and plasma CA QTL overlap on chr 7 at 122 Mbp. Likewise, four BA QTL that are all derivatives of the secondary BA DCA, including plasma TDCA and cecal DCA, isodeoxycholic acid (IDCA), and HDCA overlap on chr 12 between ∼99 – 104 Mbp. For the cecal BA, the WSB founder haplotype was associated with higher levels of these three BA, while the NOD founder haplotype was associated with lower levels. The opposite pattern was observed for plasma TDCA, where the NOD and WSB haplotype were associated with higher and lower levels, respectively (S3A-S3D Fig).

We also identified overlapping QTLs on chr 11 at ∼71 Mbp for cecal levels of the secondary BAs lithocholic acid (LCA) and isolithocholic acid (ILCA), the isomer of LCA produced by bacterial 3α-hydroxylation (S3E Fig). Higher levels of these cecal BAs are associated with the 129 founder haplotype and lower levels are associated with the A/J founder haplotype (S3F-S3G Fig). We identified the positional candidate gene *Slc13a5* (S3H Fig), which is a sodium-dependent transporter that mediates cellular uptake of citrate, an important precursor in the biosynthesis of fatty acids and cholesterol (54). Recent evidence indicates that *Slc13a5* influences host metabolism and energy homeostasis (55–57). *Slc13a5* is a transcriptional target of pregnane X receptor (PXR) (58), which also regulates the expression of genes involved in the biosynthesis, transport, and metabolism of BAs (59).

### Co-mapping analyses identifies novel interactions between bacterial taxa and bile acid homeostasis

We searched for regions of the chromosome that were associated with both BA and bacterial abundance, as this may provide evidence of interactions between the traits (60). We identified 17 instances of overlapping microbial and BA QTL on 12 chromosomes. This QTL overlap indicates there might be QTL with pleiotropic effects on BAs and the microbiota, suggest that genetic variation influencing host BA profiles has an effect on compositional features of the gut microbiota, or genetic-driven variation in microbiota composition alters BAs. Examples of notable instances of overlapping bacterial and BA QTL are discussed in the Supporting Information (S1 Data).

We focused our co-mapping analysis on chr 8 at ∼ 5.5 Mbp, where *Turicibacter sp.* QTL and plasma cholic acid (CA) QTL overlap (Fig 3A and 3B). These traits were particularly interesting because both have been shown to be influenced by host genetics by previous studies. *Turicibacter* has been identified as highly heritable in both mouse and human genetic studies (24,27,45,49), whereas multiple reports have found differences in CA levels as a function of host genotype (18,61). Furthermore, CA levels are influenced by both host genetics and microbial metabolism since it is synthesized by host liver enzymes from cholesterol and subsequently modified by gut microbes in the intestine. Notably, these co-mapping traits also share the same allele effects pattern, where the A/J and WSB haplotypes have strong positive and negative associations, respectively (Fig 3C and 3D).

**Figure 3.**
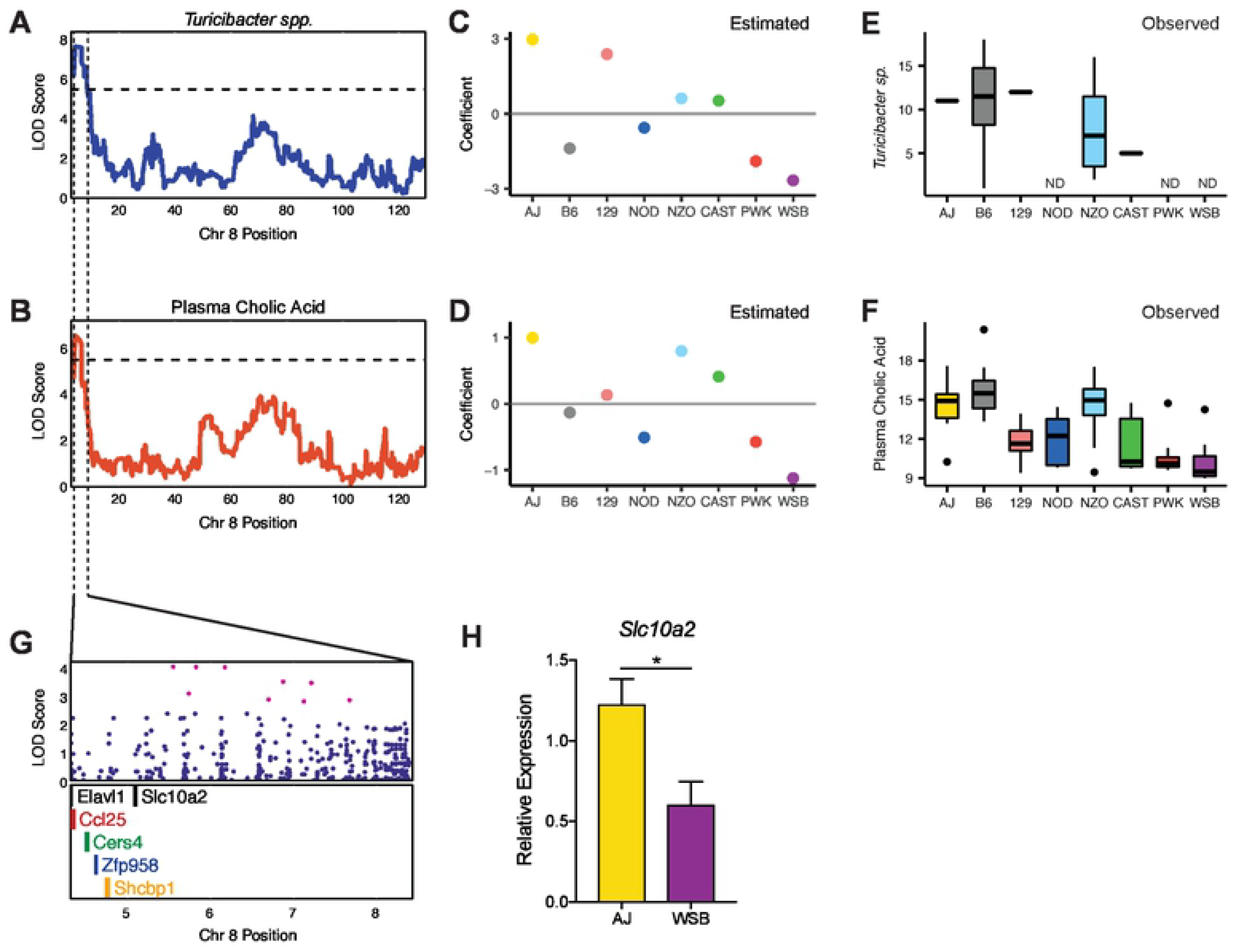
Co-mapping of *Turicibacter sp*. and plasma cholic acid (CA) QTL on chromosome 8. Association of (A) fecal abundance of *Turicibacter sp.* and (B) plasma CA levels on chromosome (chr) 8. The x-axis indicates the position in Mbp along chr 8. The y-axis for the top panel and the y-axis in the bottom panel is the LOD score. A/J and WSB founder alleles are associated with higher and lower levels of *Turicibacter* and plasma CA levels, respectively. The estimated founder strain abundance of (C) *Turicibacter* and (D) levels of plasma CA in the DO population reflects measured values observed in founder strains for (E) the abundance of *Turicibacter sp.* and (F) plasma cholic acid levels (*n* = 8 mice/genotype, 4 male and 4 female). (E) Spearman rank correlation between *Turicibacter sp*. and plasma CA in DO mice (n=192). (F) Spearman rank correlation between Turicibacter genera and plasma cholic acid in DO founder strains (n = 19). (G) SNPs (top panel) and protein coding genes (bottom panel) under the QTL interval. Magenta dots correspond to SNPs with the strongest association where the LOD drop < 1.5 from the top SNP. (H) Relative expression of *Slc10a2* measured in the distal ileum by qRT-PCR in A/J and WSB parental strains (*n* = 6, 3 male and 3 female). Data are presented as mean ± SEM; Welch’s *t* test; * p < 0.05. Correlation p-values adjusted for multiple tests using Benjamini and Hochberg correction. ND – not detected.

To assess whether the trait patterns observed in the DO founder strains correspond to the observed allelic effects in the QTL mapping, we performed a separate characterization of the fecal microbiota composition and plasma bile acids in age-matched A/J and WSB animals fed the HF/HS diet. The founder strain allele patterns inferred from the QTL mapping closely resembled the observed levels of *Turicibacter sp.* (Fig 3E) and plasma CA in the founder strains (Fig 3F), where A/J animals had significantly higher levels of *Turicibacter sp.* and CA than WSB animals. However, *Turicibacter* levels in the founder strains do not complete mirror the estimated allele effects. This may be due to other genetic factors that also influence *Turicibacter* levels, as this taxa may be influenced by multiple host genes and levels of *Turicibacter* have previously been associated on chr 7 (24), 9 and 11 (49). Furthermore, *Turicibacter* and plasma CA were positively correlated in the DO mice (r = 0.43, p = 3.53e^−10^). This finding is consistent with a previous study that found positive correlations between *Turicibacter* and unconjugated cecal BAs (62). Taken together, the overlap between the *Turicibacter sp.* QTL and plasma CA QTL, along with the similar allele effects pattern, which reflect the values observed in the founder strains, provide strong evidence suggesting that these traits are related and they are responding to the common genetic driver.

### *Slc10a2* is a candidate gene for *Turicibacter sp.* and plasma cholic acid

We searched in the QTL confidence interval for candidate genes via high-resolution association mapping on chr 8 and identified SNPs associated with both traits. Among these we identified SNPs upstream of the gene *Slc10a2*, which encodes for the apical sodium-bile transporter (Fig 3G). Slc10a2 is responsible for ∼95% of BA reabsorption in the distal ileum and plays a key role in BA homeostasis (63). In humans, mutations in this gene are responsible for primary BA malabsorption, resulting in interruption of enterohepatic circulation of BAs and decreased plasma cholesterol levels (64). Likewise, *Slc10a2*^−/−^ mice have a reduced total BA pool size, increased fecal BA concentrations and reduced total plasma cholesterol in comparison to wild-type mice (63). Additionally, a comparison between germ-free and conventionally-raised mice found that expression of *Slc10a2* is downregulated in presence of the gut microbiota, suggesting microbes may influence the expression of the transporter (41).

Our analysis identified SNPs associated with levels of *Turicibacter sp.* and plasma CA at the QTL peak (Fig 3G). The SNPs with the strongest associations were attributed to the WSB and A/J haplotypes and fell on intergenic regions near *Slc10a2*. There is growing evidence that non-coding intergenic SNPs are often located in or closely linked to regulatory regions, suggesting that they may influence host regulatory elements and alter gene expression (65,66). To assess if candidate gene expression patterns in the DO founders corresponds to the estimated allelic effects in the QTL mapping, we quantified *Slc10a2* expression in distal ileum samples from A/J and WSB mice by quantitative reverse transcriptase PCR (qRT-PCR). A/J mice exhibited significantly higher expression of *Slc10a2* compared to WSB mice (Fig 3H), which is consistent with estimated allele patterns for the overlapping *Turicibacter* and plasma CA QTLs on chr 8 (Fig 3A and 3B). Remarkably, several studies have noted concomitant changes in microbiota composition and *Slc10a2* mRNA levels (67–69).

### A common genetic driver controls *Turicibacter sp.* and plasma cholic acid

We mapped QTL for *Turicibacter sp.* and for plasma CA levels to a common locus on chr 8 at 5-7 Mbp. Since the LOD profiles and allelic effects are highly similar, the QTL may be due to a single shared locus (pleiotropy) or multiple closely linked loci. We examined this question using a likelihood ratio testing of the null hypothesis of pleiotropy versus the alternative of two independent genetic regulators of these traits (70). Analysis of 1000 bootstrap samples resulted in a p-value of 0.531, which is consistent with the presence of a single pleiotropic locus that affects both traits.

We next sought to understand the causal relationships between the microbe and the BA. We asked whether the relationship between the microbe and BA was causal, reactive or independent. To establish the directionality of the relationship, we applied mediation analysis where we conditioned one trait on the other (71). When we conditioned *Turicibacter sp.* on plasma CA (QTL → BA → Microbe), we observed a LOD drop of 3.2 (Fig 4A and 4B). Likewise, when we conditioned the plasma cholic acid on the microbe (QTL → Microbe → BA) there was a LOD drop of 3.32 (Fig 4C and 4D). The partial mediation seen in both models suggests that the relationship between the microbe and the BA could be bidirectional, where they exert an effect on one another.

**Figure 4.**
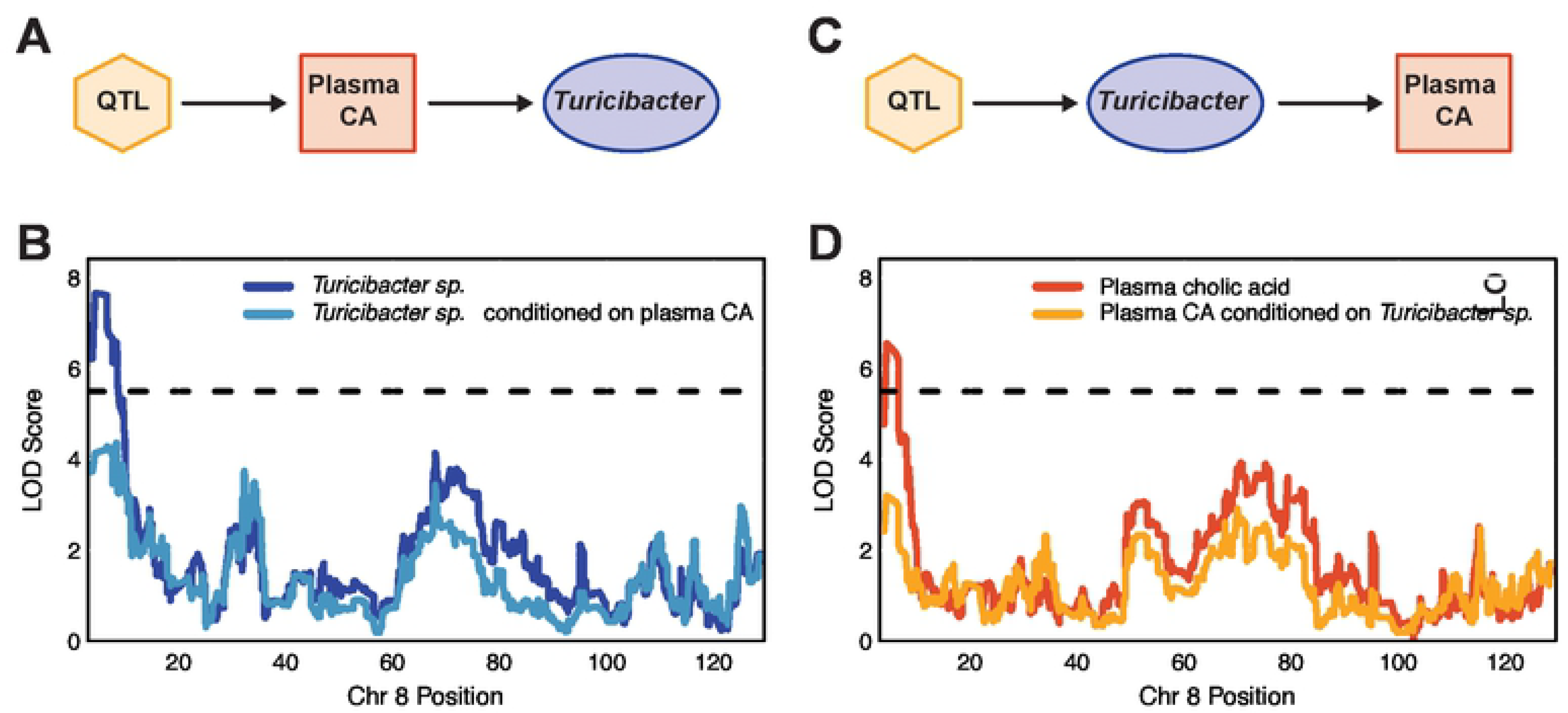
Mediation analysis and causal inference testing suggest causal relationship between *Turicibacter sp.* abundance and plasma cholic acid (CA) levels. (A) Hypothetical causal model that proposes that cholic acid (CA) mediates the changes in *Turicibacter sp.* abundance. (B) Change in LOD score of plasma CA when adjusting for *Turicibacter sp.* abundance. The x-axis indicates the position in Mbp along chr 8. (C) Hypothetical causal model that proposes that *Turicibacter sp.* mediates changes in abundance of plasma CA levels. (D) Change in LOD score of *Turicibacter sp.* when controlling for plasma CA levels.

From this analysis, we can hypothesize this relationship can be explained by a pleiotropic model, where a single locus influences a microbial and a BA trait, and the microbial trait is also reactive to changes in the BA trait. It is important to note that statistical inference only partially explains the relationship between the traits and there may be other hidden variables that may further explain the relationship. The complex relationship depicted by the causal inference testing is consistent with the interplay between gut microbes and BAs in the intestine and their known ability to influence the other.

### Bile acids inhibit *Turicibacter sanguinis* growth at physiologically relevant concentrations

Due to the strong correlative relationship between the QTL, we tested whether there was a direct interaction between bile acids and *Turicibacter*. *Turicibacter* inhabits the small intestine where BAs are secreted upon consumption of a meal (73,74). We screened the human isolate *Turicibacter sanguinis* for deconjugation and transformation activity *in vitro* by HPLC/MS-MS. We found that *T. sanguinis* deconjugated ∼96-100% of taurocholic acid and glycochenodeoxycholic acid (Fig 5A) within 24 hours. It also transformed ∼6 and 8 % of CA and CDCA to 7-dHCA and 7-ketolithocholic acid (7-KLCA), respectively (Fig 5B and 5C). The percent transformed did not increase after 24 hours (data not shown). Both of these transformations require the action of the bacterial 7α-hydroxysteroid dehydrogenase.

**Figure 5.**
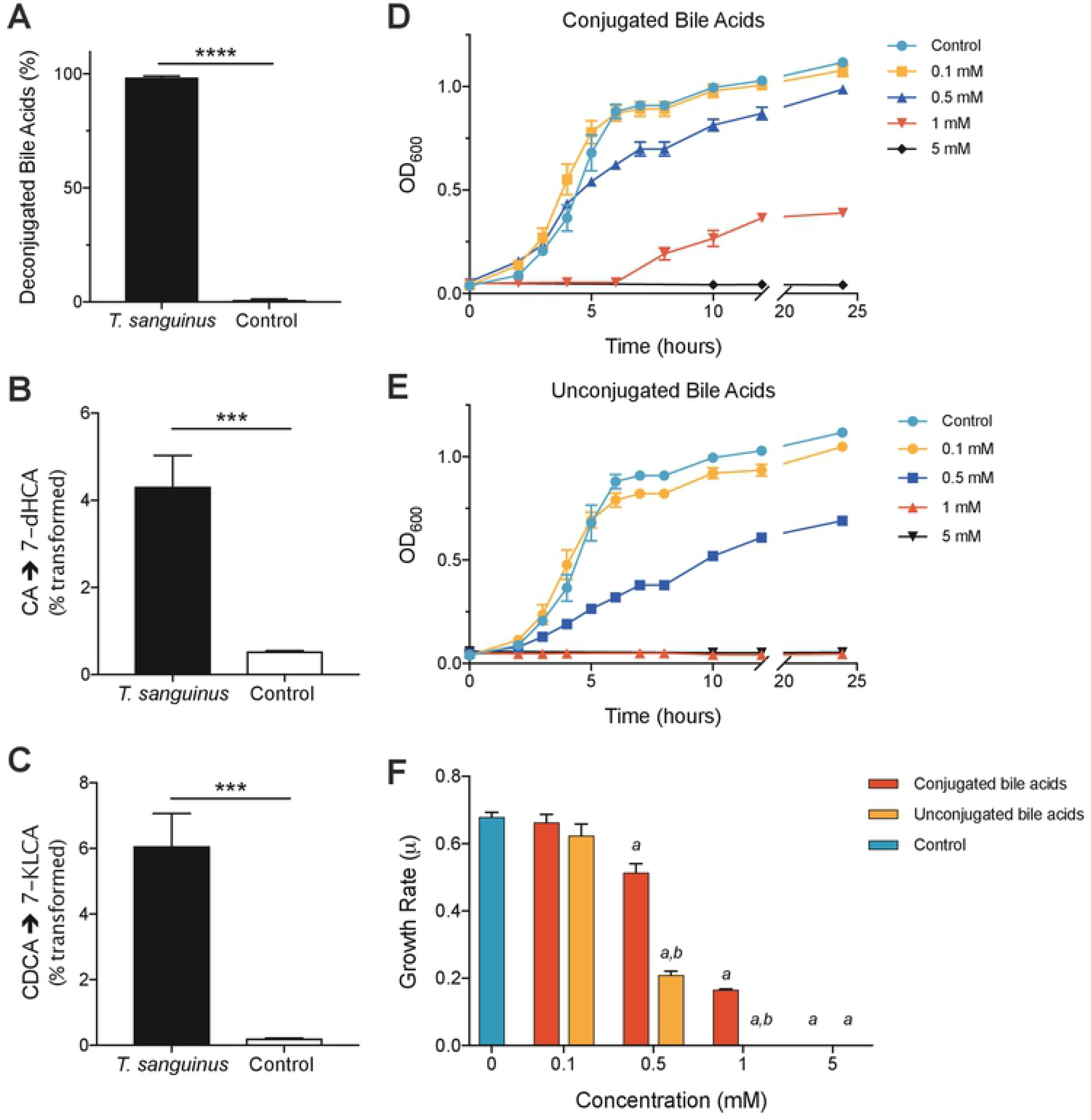
*Turicibacter sanguinis* and bile acid interactions. (A) Percent of conjugated bile acids detected after 24-hour incubation with or without the presence of *T. sanguinis*. (B) Transformation of cholic acid (CA) to 7-dehydrocholic acid (7-dHCA), and (C) chenodeoxycholic acid (CDCA) to 7-ketolithocholic acid (7-KLCA) by *T. sanguinis* after 24 hours. Growth of *T. sanguinis* in the presence of 0.1 mM, 0.5 mM, 1 mM and 5 mM (D) conjugated (equimolar pool of taurocholic acid (TCA) and glycochenodeoxycholic acid (GCDCA)), and (E) unconjugated (equimolar pool of cholic CA, CDCA, and deoxycholic acid (DCA)) bile acids over 24 hours. (F) Growth rate (μ) of *T. sanguinis* in medium supplemented with varying concentrations of conjugated and unconjugated bile acids. Data shown are from one experiment with three technical replicates. Data are presented as mean ± SEM; one-way ANOVA followed by Tukey’s multiple comparisons test; ** p < 0.01, *** p < 0.001, **** p < 0.0001.

Based on these results, we asked if conjugated and unconjugated bile acids differentially modulate *T. sanguinis* growth. BA concentrations range from ∼1-10 mM along the small intestine (75) to ∼0.2-1 mM in the cecum (76). Therefore, we grew *T. sanguinis* in the presence of either conjugated or unconjugated bile acids at physiologically relevant concentrations ranging from 0.1 – 5 mM. *T. sanguinis* growth decreased with increasing concentrations of BAs and growth was completely inhibited at 1 mM for unconjugated BAs and 5 mM for conjugated BAs (Fig 5D and 5E). Growth rate was significantly slower in the presence of 1 mM conjugated and 0.5mM unconjugated bile acids (Fig 5F). These results suggest that levels of BAs may affect abundance of *Turicibacter* in the gut.

To compare *T. sanguinis* sensitivity to conjugated bile acids relative to other small intestine colonizers, we grew four taxa (*Bacteroides thetaiotaomicron*, *Clostridium asparagaiforme*, *Lactobacillus reuteri* and *Escherichia coli* MS200-1) known to colonize this region of the intestine with or without 1 mM conjugated bile acids. Members of these genera are known to have bile salt hydrolase (BSH) activity to deconjugate bile acids (35). Unlike *T. sanguinis*, the addition of high levels of conjugated bile acids had little to no effect on the growth of these four gut microbes (S4 Fig). Consistent with these findings, *Turicibacter* abundance was negatively correlated with cecal TCA levels in the DO mice (r = −0.262, p = 0.0035).

Taken together, these data indicate that *T. sanguinis* is sensitive to higher concentrations of BA compared to other small intestine colonizers. These reciprocal effects between the BA and the bacterium provide biological evidence for the correlative relationship shown by the causal model testing. In summary, using a genetic approach, we identified and provide validation of a relationship between a genetic locus containing the BA transporter *Slc10a2*, and levels of *Turicibacter* and plasma cholic acid. Based on our findings, we hypothesize that the identified locus regulates expression of *Slc10a2*, altering active BA reabsorption in the ileum, leading to increased intestinal BA concentrations and alterations in the intestinal BA environment. Consequently, the resulting environmental change provides an unfavorable habitat for *Turicibacter.* In turn, lower levels of *Turicibacter* BA deconjugation activity leads to a decrease in circulating free plasma cholic acid levels.

## Conclusion

In this study, we performed the first known genetic mapping integration of gut microbiome and BA profiles. Using DO mice, we identified multiple QTL for gut microbes and bile acids spanning the host genome. These included loci that associated with individual microbial and BA traits, as well as loci with potential pleiotropic effects, where a single genetic region influenced both the abundance of a gut microbe and levels of a BA. While several studies suggest that host genetic variation has a minor impact on microbiota composition, there are overlapping findings among different studies in both human and mouse populations that indicate that specific bacterial taxa are influenced by host genetics. Our results in the DO population corroborate several of these key findings (discussed in S1 Data). *Turicibacter sp*. is among the microbes consistently associated with host genetics. This work plus data from previous reports suggest that alterations in the BA pool driven by *Slc10a2* genetic variation and concomitant changes in expression/activity elicit an impact on gut microbiota community structure and influence the ability of *Turicibacter* to colonize and persist in the intestine. Although this microbe deconjugates primary BAs, we found that it is also sensitive to elevated concentrations of both conjugated and unconjugated BAs. Future experiments are needed to examine how a decrease in *Slc10a2* expression changes intestinal BA profiles and the consequences on *Turicibacter* colonization. Additionally, this work identified multiple host-microbe-metabolite interactions that need to be validated with additional molecular studies. More broadly, our work demonstrates the power of genetics to identify novel interactions between microbial and metabolite traits and provides new testable hypotheses to further dissect factors that shape gut microbiota composition.

## Materials and methods

### Animals and sample collection

Animal care and study protocols were approved by the University of Wisconsin-Madison Animal Care and Use Committee. DO mice were obtained from the Jackson Laboratories (Bar Harbor, ME, USA) at ∼4 weeks of age and maintained in the Department of Biochemistry vivarium at the University of Wisconsin-Madison. Mice were housed on a 12-hour light:dark cycle under temperature- and humidity-controlled conditions. Five waves of 100 DO mice each from generations, 17, 18, 19, 21, and 23 were obtained at intervals of 3-6 months. Each wave was composed of equal numbers of male and female mice. All mice were fed a high-fat high-sucrose diet (TD.08811, Envigo Teklad, 44.6% kcal fat, 34% carbohydrate, and 17.3% protein) *ad libitum* upon arrival to the facility. Mice were kept in the same vivarium room and were individually housed to monitor food intake and prevent coprophagy between animals. DO mice were sacrificed at 22-25 weeks of age.

The eight DO founder strains (C57BL/6J, A/J, 129S1/SvImJ, NOD/ShiLtJ, NZO/HILtJ, PWK/PhJ, WSB/EiJ and CAST/EiJ) were obtained from the Jackson Laboratories. Mice were bred at the University of Wisconsin-Madison Biochemistry Department. Mice were housed by strain and sex (2-5 mice/cage), with the exception of CAST that required individual housing. Inbred founder mice were housed under the same environmental conditions as the DO animals. Like the DO mice, the eight founder strains were maintained on the HF/HS diet and were sacrificed at 22 weeks of age, except for NZO males that were sacrificed at 14 weeks, due to high mortality attributable to severe disease.

For both DO and founder mice, fecal samples for 16S rRNA sequencing were collected immediately before sacrifice after a 4 hour fast. Cecal contents, plasma, and additional tissues were harvested promptly after sacrifice and all samples were immediately flash frozen in liquid nitrogen and stored at −80°C until further processing.

### DNA extraction

DNA was isolated from feces using a bead-beating protocol {Turnbaugh:2009ei}. Mouse feces (∼1 pellet per animal) were re-suspended in a solution containing 500μl of extraction buffer [200mM Tris (pH 8.0), 200mM NaCl, 20mM EDTA], 210μl of 20% SDS, 500μl phenol:chloroform:isoamyl alcohol (pH 7.9, 25:24:1) and 500μl of 0.1-mm diameter zirconia/silica beads. Cells were mechanically disrupted using a bead beater (BioSpec Products, Barlesville, OK; maximum setting for 3 min at room temperature), followed by extraction with phenol:chloroform:isoamyl alcohol and precipitation with isopropanol. Contaminants were removed using QIAquick 96-well PCR Purification Kit (Qiagen, Germantown, MD, USA). Isolated DNA was eluted in 5 mM Tris/HCL (pH 8.5) and was stored at −80°C until further use.

### 16S rRNA Sequencing

PCR was performed using universal primers flanking the variable 4 (V4) region of the bacterial 16S rRNA gene (102). Genomic DNA samples were amplified in duplicate. Each reaction contained 10-30 ng genomic DNA, 10 µM each primer, 12.5 µl 2x HiFi HotStart ReadyMix (KAPA Biosystems, Wilmington, MA, USA), and water to a final reaction volume of 25 µl. PCR was carried out under the following conditions: initial denaturation for 3 min at 95°C, followed by 25 cycles of denaturation for 30 s at 95°C, annealing for 30 s at 55°C and elongation for 30 s at 72°C, and a final elongation step for 5 min at 72°C. PCR products were purified with the QIAquick 96-well PCR Purification Kit (Qiagen, Germantown, MD, USA) and quantified using Qubit dsDNA HS Assay kit (Invitrogen, Oregon, USA). Samples were equimolar pooled and sequenced by the University of Wisconsin – Madison Biotechnology Center with the MiSeq 2×250 v2 kit (Illumina, San Diego, CA, USA) using custom sequencing primers.

### 16S analysis

Demultiplexed paired end fastq files generated by CASAVA (Illumina) and a mapping file were used as input files. Sequences were processed, quality filtered and analyzed with QIIME2 (version 2018.4) (https://qiime2.org), a plugin-based microbiome analysis platform (103). DADA2 (47) was used to denoise sequencing reads with the q2-dada2 plugin for quality filtering and identification of *de novo* exact sequence variants (ESVs) (i.e. 100% exact sequence match). This resulted in 20,831,573 total sequences with an average of 52,078 sequences per sample for the DO mice, and 2,128,796 total sequences with an average of 34,335.4 sequences per sample for the eight DO founder strains. Sequence variants were aligned with mafft (104) with the q2-alignment plugin. The q2-phylogeny plugin was used for phylogenetic reconstruction via FastTree (105). Taxonomic classification was assigned using classify-sklearn (106) against the Greengenes 13_8 99% reference sequences (107). Alpha- and beta-diversity (weighted and unweighted UniFrac (108) analyses were performed using q2-diversity plugin at a rarefaction depth of 10000 sequences per sample. For the DO mice, one sample (DO071) was removed from subsequent analysis because it did not reach this sequencing depth. For analysis of the eight DO founder strains, one sample (NOD5) was removed because it did not reach this sequencing depth. Subsequent processing and analysis were performed in R (v.3.5.1), and data generated in QIIME2 was imported into R using Phyloseq (109). Sequencing data was normalized by cumulative sum scaling (CSS) using MetagenomeSeq (110). Summaries of the taxonomic distributions were generated by collapsing normalized ESV counts into higher taxonomic levels (genus to phylum) by phylogeny. We defined a core measurable microbiota (CMM) (24) to include only microbial traits present in 20% of individuals in the QTL mapping. In total, 86 ESVs and 42 collapsed microbial taxonomies comprised the CMM.

### Sample preparation for plasma bile acid analysis

40 μL of DO plasma collected at sacrifice (30 μL used for founder strains) were aliquoted into a tube with 10 μL SPLASH Lipidomix internal standard mixture (Avanti Polar Lipids, Inc.). Protein was precipitated by addition of 215 μL MeOH. After the mixture was vortexed for 10 s, 750 μL methyl tert-butyl ether (MTBE) were added as extraction solvent and the mixture was vortexed for 10 s and mixed on an orbital shaker for 6 min. Phase separation was induced by adding 187.5 μL of water followed by 20 s of vortexing. All steps were performed at 4 °C on ice. Finally, the mixture was centrifuged for 4 min at 14,000 x g at 4 °C and stored at −80 °C. For targeted bile acids analysis, samples were thawed on ice. 400 μL of ethanol were added to further precipitate protein, as well as 15 μL of isotope-labeled internal standard mix (12.5 µM d4-TαMCA, 10 µM d4-CDCA). The samples were vortexed for 20 s and centrifuged for 4 min at 14,000 g at 4 °C after which the supernatant (ca. 1000 μL) was taken out and dried down. Dried supernatants were resuspended in 60 μL mobile phase (50 %B), vortexed for 20 s, centrifuged for 4 min at 14,000 g and then 50 μL were transferred to vials with glass inserts for MS analysis.

### Sample preparation for cecal bile acid analysis

30 ± 7.5 mg cecal contents along with 10 μL SPLASH Lipidomix internal standard mixture were aliquoted into a tube with a metal bead and 270 μL MeOH were added for protein precipitation. To each tube, 900 μL MTBE and 225 μL of water were added as extraction solvents. All steps were performed at 4 °C on ice. The mixture was homogenized by bead beating for 8 min at 25 Hz. Finally, the mixture was centrifuged for 4-8 min at 11,000 x g at 4 °C. Subsequent processing for the DO mice and eight DO founder strains differed due to other analyses performed on the samples that are not presented in this paper. For DO samples, 100 μL of the aqueous and 720 μL of organic layer were combined and stored at −80 °C. For analysis, these were thawed on ice and 400 μL of ethanol were added to further precipitate protein, as well as 15 μL of isotope-labeled internal standard mix (12.5 µM d4-TαMCA, 10 µM d4-CDCA). The samples were vortexed for 20 s and centrifuged for 4 min at 14,000 g at 4 °C after which the supernatant (ca. 1000 μL) was taken out and dried down. Dried supernatants were resuspended in 100 μL mobile phase (50 %B), vortexed for 20 s, centrifuged for 8 min at 14,000 g and then 50 μL were transferred to vials with glass inserts for MS analysis. For the eight DO founder strains, the mixture was dried down including all solid parts and stored dried at −80 °C. For targeted bile acid analysis, these dried down samples were then thawed on ice and reconstituted in 270 μL of methanol, 900 μL of MTBE, and 225 μL of water. 400 μL of ethanol were added to further precipitate protein, as well as 15 μL of isotope-labeled internal standard mix (12.5 µM d4-TαMCA, 10 µM d4-CDCA). The mixture was bead beat for 8 min at 25 Hz and centrifuged at 14,000 g for 8 minutes after which the supernatant (ca. 1500 μL) was taken out and dried down. Dried supernatants were resuspended in 100 μL mobile phase (50 %B), vortexed for 20 s, centrifuged for 4 min at 14,000 g and then 90 μL were transferred to vials with glass inserts for MS analysis.

### Measurement and analysis of mouse bile acids

LC-MS analysis was performed in randomized order using an Acquity CSH C18 column held at 50 °C (100 mm × 2.1 mm × 1.7 μm particle size; Waters) connected to an Ultimate 3000 Binary Pump (400 μL/min flow rate; Thermo Scientific). Mobile phase A consisted of 10 mM ammonium acetate containing 1 mL/L ammonium hydroxide. Mobile phase B consisted of MeOH with the same additives (111). Mobile phase B was initially held at 50% for 1.5 min and then increased to 70% over 13.5 min. Mobile phase B was further increased to 99% over 0.5 min and held for 2.5 min. The column was re-equilibrated for 5.5 min before the next injection. Twenty microliters of plasma sample or ten microliters of cecum sample were injected by an Ultimate 3000 autosampler (Thermo Scientific). The LC system was coupled to a TSQ Quantiva Triple Quadrupole mass spectrometer (Thermo Scientific) by a heated ESI source kept at 325°C (Thermo Scientific). The inlet capillary was kept at 350 °C, sheath gas was set to 15 units, auxiliary gas to 10 units, and the negative spray voltage was set to 2,500 V. For targeted analysis the MS was operated in negative single reaction monitoring (SRM) mode acquiring scheduled, targeted scans to quantify selected bile acid transitions, with two transitions for each species’ precursor and 3 min retention time windows. Collision energies were optimized for each species and ranging from 20-55 V. Due to insufficient fragmentation for unconjugated bile acids, the precursor was monitored as one transition with a CE of 20 V. MS acquisition parameters were 0.7 FWHM resolution for Q1 and Q3, 1 s cycle time, 1.5 mTorr CID gas and 3 s Chrom filter. In total, 27 bile acids, including 14 unconjugated, 9 tauro- and 4 glycine-conjugated species, were measured. The resulting bile acid data were processed using Skyline 3.6.0.10493 (University of Washington). For each species, one transition was picked for quantitation, while the other was used for retention time confirmation. Normalization of the quantitative data was performed to the internal standard d4-CDCA as indicated in Equation 1.

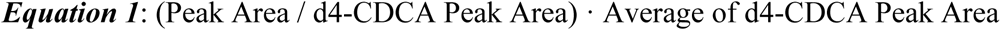

### Genotyping

Genotyping was performed on tail biopsies as previously described (42) using the Mouse Universal Genotyping Array (GigaMUGA) [143,259 markers] (112) at Neogen (Lincoln, NE). Genotypes were converted to founder strain-haplotype reconstructions using a hidden Markov model (HMM) implemented in the R/qtl2 package (48). We interpolated the GigaMUGA markers onto an evenly spaced grid with 0.02-cM spacing and added markers to fill in regions with sparse physical representation, which resulted in 69,005 pseudomarkers.

### QTL mapping

We performed QTL mapping using the R package R/qtl2 (48). QTL mapping was done through a regression of the phenotype on the founder haplotype probabilities estimated with an HMM designed for multi-parental populations. Genome scans were performed for each phenotype with sex, cohort (wave), and days on diet included as additive covariates. Genetic similarity between mice was accounted for using a kinship matrix based on the leave-one-chromosome-out (LOCO) methods (113). For microbial QTL mapping, normalized gut microbiota abundance data transformed to normal quantiles. For bile acid QTL mapping, normalized plasma and cecal bile acid levels were log2 transformed. The mapping statistic reported is log of the odds ratio (LOD). The significance thresholds were determined by performing 1000 permutations of genome-wide scans by shuffling phenotypic data in relation to individual genotypes. Significant QTL were determined at a genome-wide *P*-value of < 0.05 and the QTL support interval was defined using the 95% Bayesian confidence interval.

### Mediation/Pleiotropy analysis

To assess whether two co-mapping traits were caused by a pleiotropic locus, we used a likelihood ratio test implemented with the open source R package R/qtl2pleio (70). Here, we compared the alternative hypothesis of two distinct loci with the null hypothesis of pleiotropy for two traits that map to the same genetic region. Parametric bootstrapping was used to determine statistical significance. Mediation analysis was applied to identify whether a microbe or bile acid were likely to be a causal mediator of the QTL as presented in Li et al. (114). This analysis was adapted from a general approach previously described to differentiate target from mediator variables (115). The effect of a mediator on a target was evaluated by performing an allele scan or SNP scan using the target adjusted by mediator. Only individuals with both values for both traits were considered for mediation analysis. Traits with a LOD drop >2 after controlling for the mediator were considered for further causality testing. To statistically assess causality between microbial and bile acid trait sets (causal, reactive, independent, undecided), a causal model selection test (72) was applied using the R packages R/intermediate and R/qtl2. Causal model selection tests were evaluated on both alleles and SNPs in peak region.

### RNA extraction

Total RNA was extracted from flash-frozen distal ileum tissues by TRIzol extraction and further cleaned using the RNeasy Mini Kit (Qiagen, Germantown, MD, USA). DNA was removed by on-column DNase digestion (Qiagen). Purified RNA was quantified using a Nanodrop 2000 spectrophotometer.

### Quantitative Real-Time PCR

SuperScript II Reverse Transcriptase with oligo(dT) primer (all from Invitrogen, Carlsbad, CA, USA) was used to synthesize 20 μl cDNA templates from 1 μg purified RNA. cDNA was diluted 2X before use and qRT-PCR reactions were prepared in a 10 μl volume using SsoAdvanced Universal SYBR Green Supermix (Bio-Rad, Hercules, CA, USA) and 400 nM specific primers targeting the gene of interest (SLC10A2-F [5’-TGGGTTTCTTCCTGGCTAGACT-3’]; SLC10A2-R [5’-TGTTCTGCATTCCAGTTTCCAA-3’] (116)). All reactions were performed in triplicate. Reactions were run on a CFX96 Real-Time PCR System (Bio-Rad, Hercules, CA, USA). The 2^−∆∆Ct^ method (117) was used to calculate relative changes in gene expression and all results were normalized to GAPDH.

### Bacterial culturing

Bacterial strains were obtained from DSMZ and ATCC. All strains were cultured at 37°C under anaerobic conditions using an anaerobic chamber (Coy Laboratory Products) with a gas mix of 5% hydrogen, 20% carbon dioxide and 75% nitrogen. Strains were grown in rich medium (S5 Table) that was filter sterilized and stored in the anaerobic chamber at least 24 hours prior to use. *L. reuteri* was grown in medium supplemented with 20 mM glucose. For all *in vitro* assays, cultures used for inoculation were grown overnight at 37°C in 10 mL 14b medium in anaerobic Hungate tubes. Stock solutions of conjugated bile acids (TCA, GCDCA) and unconjugated bile acids (CA, CDCA, DCA) were prepared to a final concentration of 100 mM and used for all *in vitro* assays. All bile acids used were soluble in methanol.

### Microbial bile acid metabolism screen

Stock solutions of conjugated and unconjugated bile acids (100 mM) were added to 3 ml 14b medium to obtain a final concentration of 100 μM total bile acid. Tubes were inoculated with a *T. sanguinis* cultured overnight, then incubated in the anaerobic chamber at 37°C for 48 hours. At the 24- and 48-hour timepoints, 1 mL of each culture was removed and the supernatant was collected after brief centrifugation. Each culture supernant was diluted 10x in initial running solvent (30:70 MeOH:10 mM ammonium acetate). Samples were spun at max speed for 3 minutes to remove suspended particles prior to loading on the uHPLC. Samples were analyzed using a uHPLC coupled with a high-resolution mass spectrometer.

### Microbial bile acid screen uHPLC-MS/MS parameters

10 µL aliquots of diluted supernatant samples were analyzed using a uHPLC-MS/MS system consisting of a Vanquish uHPLC coupled by electrospray ionization (ESI) (negative mode) to a hybrid quadrupole-high-resolution mass spectrometer (Q Exactive Orbitrap; Thermo Scientific). Liquid chromatography separation was achieved on an Acquity UPLC BEH C_18_ column (2.1-by 100-mm column, 1.7-µm particle size) heated to 50°C. Solvent A was 10 mM Ammonium acetate, pH 6; solvent B was 100% methanol. The total run time was 31.5 minutes with the following gradient: 0 min, 30% B; 0.5 min, 30% B; 24 min, 100% B; 29 min, 100% B; 29 min, 30% B; 31.5 min, 30% B. Bile acid peaks were identified using the Metabolomics Analysis and Visualization Engine (MAVEN) (118).

### Growth curves

Bacterial growth rate was measured in medium 14b supplemented with either 100 μM, 300 μM, 1 mM bile acids or methanol control. Medium was dispensed inside an anerobic chamber into Hungate tubes. Tubes containing 10 mL of medium were inoculated with 30 μL of an overnight culture and incubated at 37°C for 24 hours. *T. sanguinis* was grown with shaking to disrupt the formation of flocculent colonies. Growth was monitored as the increase in absorbance at 600 nm in a Spectronic 20D+ spectrophotometer (Thermo Scientific, Waltham, MA, USA). Growth rate was determined as μ = ln(*X*/*X_o_*)/*T*, where *X* is the OD_600_ value during the linear portion of growth and *T* is time in hours. Values given are the mean μ values from two independent cultures done in triplicate.

### Statistical analysis

All statistical analyses were performed in R (v.3.5.1) (119). Unless otherwise indicated in the figure legends, differences between groups were evaluated using unpaired two-tailed Welch’s t-test. For multiple comparisons, Krustkal-Wallis test was used if ANOVA conditions were not met, followed by Mann-Whitney/Wilcoxon rank-sum for multiple comparisons and adjusted for multiple testing using the Benjamini-Hochberg FDR procedure. The correlation between the abundance of microbial taxa was performed using Spearman’s correlation in the “Hmisc” (v.4.1-1) R package (120). The p-values were adjusted using the Benjamini and Hochberg method, and correlation coefficients were visualized using the “pheatmap” (v.1.0.10) (121). Multiple groups were compared by Kruskal-Wallis test and adjusted for multiple testing using the Benjamini-Hochberg FDR procedure. Significance was determined as p-value < 0.05. To assess magnitude of variability of the CMMs, summary statistics were calculated on each CMM (taxa and ESVs). Non-parametric-based PERMANOVA statistical test (122) with 999 Monte Carlo permutations was used to compare microbiota compositions among groups using the Vegan R package (123).

## Acknowledgements

The authors thank the University of Wisconsin Biotechnology Center DNA Sequencing Facility for providing sequencing and support services, and the University of Wisconsin Center for High Throughput Computing (CHTC) in the Department of Computer Sciences for providing computational resources, support, and assistance. We also thank Paul Dawson for his feedback.

## Supporting information legends

**S1 Figure. Principal coordinate analysis (PCoA) of unweighted UniFrac distances for fecal samples.** PCoA shows significant clustering by (A) sex (F = 5.572, p = 0.001) and (B) wave (F = 16.954, p = 0.001). Clustering by treatment evaluated by PERMANOVA.

**S2 Figure. Plasma and cecal bile acids group by sex, but not wave.** PCAs of plasma bile acid profiles colored by (A) sex (p < 0.0001) and (B) wave (p = 0.594), and PCAs of cecal bile acid profiles colored by (C) sex (p = 0.011) and (D) wave (p = 0.207). Kruskal Wallis one-way test followed by Wilcoxon pair-wise multiple comparisons with Benjamini and Hochberg correction.

**S3 Figure. Related bile acid species map associate to same locus.** (A) Haplotype effects and LOD scores of plasma taurodeoxycholic acid (TDCA), (B) cecal deoxycholic acid (DCA), (C) cecal isodeoxycholic acid (IDCA) and (D) cecal hyodeoxycholic acid (HDCA). For each plot, the x-axis is the physical position in Mbp along chr 12. The y-axis for the top panel is the effect coefficient depicting the estimated contributions of each founder allele, and the y-axis in the bottom panel is the LOD score. (E) Cecal levels of isolithocholic acid (ILCA) and lithocholic acid (LCA) associate to same locus on chr 11. (F) Estimated founder allele effects for cecal ILCA and (G) LCA. (H) Genes under cecal LCA and ILCA QTL interval. Dashed lines denote QTL confidence interval.

**S4 Figure. Gut associated bacteria have differential growth responses to conjugated bile acids.** Growth rate in the presence of 1 mM conjugated bile acids or methanol control for (A) *Bacteroides thetaiotaomicron*, (B) *Clostridium asparagiforme*, (C) *Escherichia coli* MS200-1, and (D) *Lactobacillus reuteri.* Data shown are from duplicate experiments with three technical replicates. Data are presented as mean ± SEM; Welch’s *t* test; no significant differences were observed between growth conditions for any of the tested organisms.

**S5 Figure. Peptostreptococcaceae and plasma bile acids co-map on chromosome (chr) 3**. Haplotype effects and LOD scores of (A) Peptostreptococcaceae family, (B) plasma cholic acid (CA), (C) plasma chenodeoxycholic acid (CDCA), (D) plasma muricholic acid (MCA), (E) plasma ursodeoxycholic acid (UDCA), and (F) plasma 7-dehydrocholic acid (7-dHCA). For each plot, the x-axis is the physical position in Mbp along chr 3. The y-axis for the top panel is the effect coefficient depicting the estimated contributions of each founder allele, and the y-axis in the bottom panel is the LOD score. All overlapping QTL have positive association with the NOD allele. (G) Protein coding genes under QTL interval.

**S6 Figure. Exact sequence variant of *Akkermansia muciniphila* and plasma bile acid QTL overlap on chromosome (chr) 1.** Haplotype effects and LOD scores of (A) *A. muciniphila* (B) plasma cholic acid (CA), (C) plasma muricholic acid (MCA), and (D) plasma 7-dehydrocholic acid (7-dHCA). For each plot, the x-axis is the physical position in Mbp along chr 1. The y-axis for the top panel is the effect coefficient depicting the estimated contributions of each founder allele, and the y-axis in the bottom panel is the LOD score. (E) Protein coding genes under 10 Mbp QTL interval. Spearman correlations in the DO mice between *A. muiniphila* and (F) plasma CA, (G) plasma MCA, and (H) plasma 7-dHCA levels. Correlation p-values adjusted for multiple tests using Benjamini and Hochberg correction. Higher levels of these microbial and bile acid traits were associated with the NZO haplotype and lower levels were associated with the 129 haplotype. (E) Protein coding genes under 10 Mbp QTL interval. Dashed lines denote QTL confidence interval. Spearman correlations in the DO mice between *A. muiniphila* and (F) plasma CA, (G) plasma MCA, and (H) plasma 7-dHCA levels. Correlation p-values adjusted for multiple tests using Benjamini and Hochberg correction.

**S1 Table. Measures of variability of microbial exact sequence variants (ESVs) or taxon (phylum, class, order, family, genus) in DO mice**. Data presented as normalized read counts; n = 399; SD, standard deviation.

**S2 Table. Measures of variability of cecal and plasma bile acids in DO mice.** Bile acid levels are presented as log2(peak area); n = 384; SD, standard deviation.

**S3 Table. Correlations among microbial taxa, bile acid and weight traits.** Spearman’s rank correlation. Only microbial exact sequence variants, genera and family included in figure. Correlations shown passed FDR < 0.01 cut-off and correlation coefficient either < −0.35 or > 0.35. Correlating bile acids from same tissue removed from table for brevity.

**S4 Table. QTL peaks for gut microbiota, plasma and cecal bile acid, and weight traits in the Diversity Outbred mice.** Only QTL with LOD > 5.5 shown. “Pos” is peak position is Mbp. “ci_lo” and “ci_hi” correspond to the positions for the 95% bayesian confidence interval.

S5 Table. Media used for bacterial culture. Medium 14(b) recipe.

